# Convergent depression of activity-dependent bulk endocytosis in rodent models of autism spectrum disorders

**DOI:** 10.1101/2024.10.10.617607

**Authors:** Katherine Bonnycastle, Peter C. Kind, Michael A. Cousin

## Abstract

The key pathological mechanisms underlying autism spectrum disorders (ASDs) remain relatively undetermined, potentially due to the heterogenous nature of the condition. Targeted studies of a series of monogenic ASDs have revealed postsynaptic dysfunction as a central conserved mechanism. Presynaptic dysfunction is emerging as an additional disease locus in neurodevelopmental disorders, however it is unclear whether this dysfunction drives ASDs or is an adaptation to the altered brain microenvironment. To determine this, we performed a high content analysis of key stages of the synaptic vesicle lifecycle in a series of preclinical models of monogenic ASDs. These models were specifically selected to have perturbations in a diverse palette of genes that were expressed either at the pre- or post-synapse. However, all models displayed a common trait of hyperexcitability. We determined that SV fusion events and SV cargo trafficking were unaffected in all models investigated. However, a key convergent phenotype was revealed, a depression in activity-dependent bulk endocytosis (ADBE). This suggests that the depression in ADBE is a homeostatic mechanism to correct hyperexcitability in the ASD brain.

## Introduction

Autism spectrum disorders (ASDs) impact up to 1 in 150 children, with intellectual disability (ID) being a frequent co-morbidity (Mefford et al., 2012). Despite this high prevalence, the mechanisms that result in brain dysfunction at the molecular, cell and circuit level in these neurodevelopmental disorders are largely undetermined. The study of monogenic ASDs provides an opportunity to decipher these key mechanisms via the identification of a molecular locus (Krol & Feng, 2018). Interestingly, a number of the most prevalent genetic causes of these disorders concentrate on genes that control synaptic events (Yuen et al., 2017). In support, multiple preclinical monogenic ASD models display defects at the synaptic and circuit level, consistent with perturbations observed in humans with similar mutations (Contractor et al., 2021; Sahin & Sur, 2015).

The synapse is comprised of the presynapse, which releases chemical neurotransmitters during invasion of action potentials (APs), and the postsynapse which integrates this input and adapts output via a series of plastic changes. Postsynaptic alterations are commonly observed in rodent models of ASDs, supporting the view that this subcellular structure is a physical convergence point in the genesis and expression of ASDs (Bonsi et al., 2022). Key proteins include synaptic Ras GTPase-activating protein 1 (SynGAP1) and fragile X messenger ribonucleoprotein (FMRP), which control synaptic plasticity via excitatory AMPA receptor trafficking and protein translation downstream from postsynaptic metabotropic receptors respectively (Bear et al., 2004; Gamache et al., 2020; Sidorov et al., 2013). Furthermore, dysfunction in cell adhesion molecules such as presynaptic neurexins and postsynaptic neuroligins also commonly result in ASDs via incorrect stabilisation and maintenance of synapse function (Cao & Tabuchi, 2017; Craig & Kang, 2007; Uchigashima et al., 2021). Therefore, dysfunction in a series of genes, including those encoding key postsynaptic molecules is a highly prevalent cause of ASDs.

In contrast to the postsynapse, presynaptic dysfunction in neurodevelopmental disorders is still relatively under-researched. This in spite of a cohort of key genes essential for neurotransmitter release having been identified as causal in conditions such as epilepsy, ID and ASDs (Bonnycastle et al., 2021; Verhage & Sørensen, 2020). Neurotransmitter release occurs in response to the activity-dependent influx of calcium, via voltage-gated channels, which triggers the fusion of neurotransmitter-containing synaptic vesicles (SVs) (Südhof, 2013). The functional pool of SVs is relatively small in typical central nerve terminals, making their efficient regeneration essential for the maintenance of neurotransmission. This is mediated by a series of endocytosis modes, which are recruited in both time and space by different patterns of neuronal activity (Chanaday et al., 2019; Kononenko & Haucke, 2015). SV cargo retrieval is dependent on clathrin-mediated endocytosis, which can occur at either the presynaptic plasma membrane or on intracellular endosomes generated by either ultrafast or activity-dependent bulk endocytosis (ADBE, (Cousin, 2017; Kononenko et al., 2014; Watanabe et al., 2014). Ultrafast endocytosis saturates rapidly during AP trains (Soykan et al., 2017), whereas ADBE is the dominant endocytosis mode during intense neuronal activity (Clayton et al., 2008; Kokotos & Cousin, 2015).

We recently discovered a selective defect in ADBE in neurons lacking FMRP (Bonnycastle et al., 2022), which translated into a decrease in presynaptic performance during periods of high activity. Since this phenotype could be corrected via agonism of BK channels, we hypothesized that the depression of ADBE was an adaptation to the hyperexcitability observed in *Fmr1^-/y^* neurons and circuits (Booker et al., 2019; Das Sharma et al., 2020; Zhang et al., 2014). In this study, we tested this hypothesis by examining SV recycling in a cohort of five independent monogenic ASDs models which all display hyperexcitability. We discovered that neurons from all models displayed depressed ADBE, with no obvious deficit in SV exocytosis or cargo trafficking. This therefore supports the hypothesis that a reduction in ADBE is a specific homeostatic adaptation to altered brain circuit activity in ASD models.

## Results

### High content screening of SV recycling

To determine whether the observed depression of ADBE in *Fmr1^-/y^*neurons (Bonnycastle et al., 2022) was a wider synaptic signature of ASDs, we examined SV recycling using a battery of optical assays in primary cultures of hippocampal neurons derived from these models. A high content protocol was designed to capture a series of different SV recycling parameters using the genetically-encoded reporter synaptophysin-pHluorin (sypHy, Figure 1). SypHy consists of the abundant SV cargo protein synaptophysin, that has a pH-sensitive EGFP (pHluorin) inserted into an intralumenal loop (Granseth et al., 2006). It reports the pH of its immediate environment, with fluorescence quenched in the acidic SV interior and unquenched during SV fusion. Therefore, the extent of SV fusion can be estimated by the evoked increase in fluorescence during stimulation. SypHy is then retrieved from the plasma membrane and packaged to SVs, which are acidified to permit neurotransmitter filling. SV cargo retrieval is rate limiting when compared to acidification (Atluri & Ryan, 2006; Granseth et al., 2006) but see (Egashira et al., 2015), meaning that the kinetics and extent of the former can be estimated from the fluorescence decrease after stimulation terminates.

**Figure 1.**
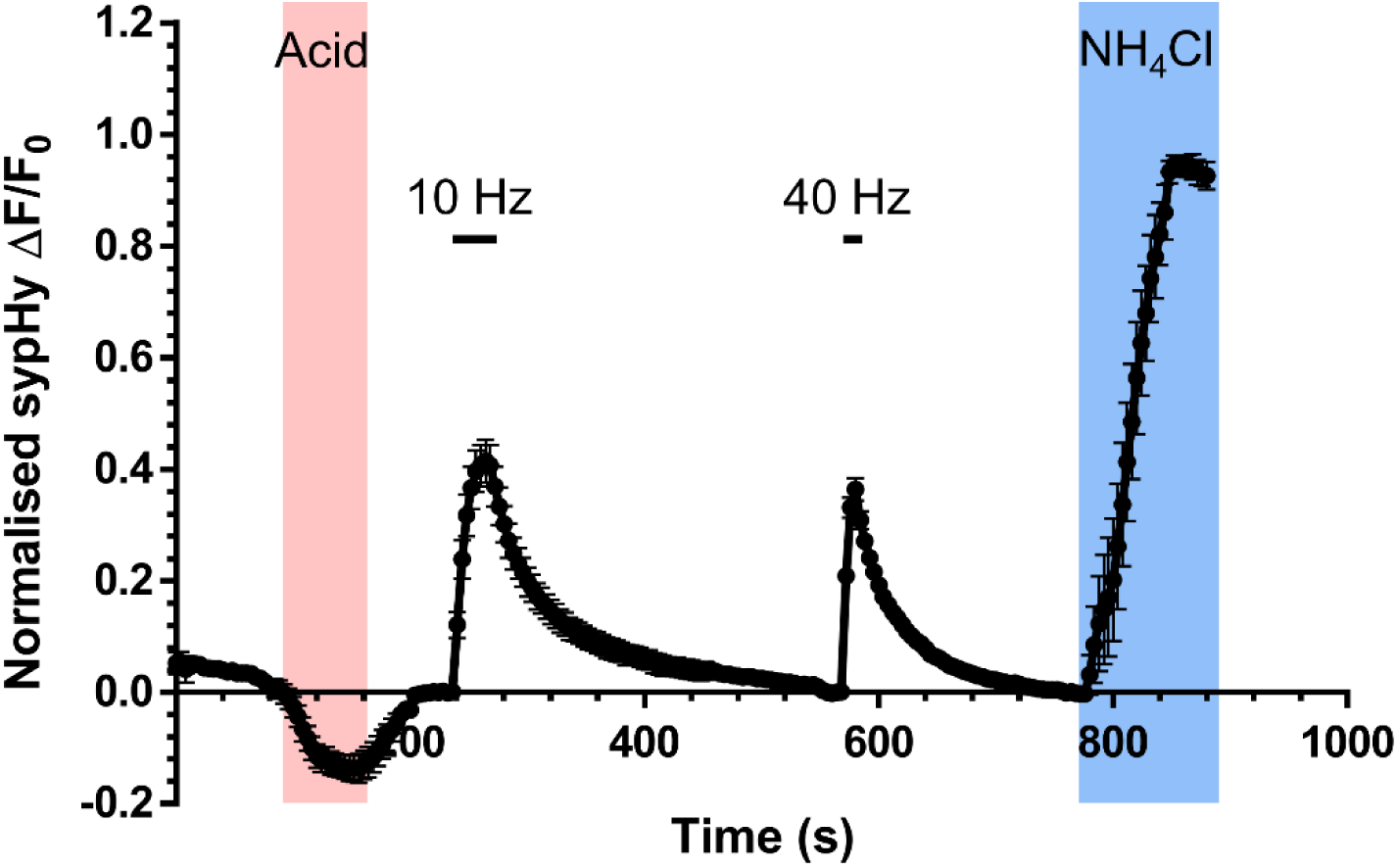
High content monitoring of SV recycling. Hippocampal neurons were transfected with synaptophysin-pHluorin (sypHy) on day *in vitro* (*DIV)* 7 and imaged at *DIV* 13-15. Transfected neurons were challenged with a pulse of impermeant acidic buffer for 1 min (to reveal the surface fraction of sypHy, shaded pink). After returning to imaging buffer for 2 min, neurons were stimulated with a train of 300 action potentials (10 Hz) followed by a 5 min rest period. After this rest period neurons were stimulated with a further train of 400 action potentials (40 Hz). Finally, after a further 3 min neurons were exposed to an alkaline buffer (NH_4_Cl, to reveal the total SV pool, blue shaded region). Stimulation is indicated by bars. Mean trace displays the average sypHy fluorescent response of wild-type neurons ± SEM. Traces are ΔF/F_0_ and are presented as a fraction of the total SV pool.

First, sypHy-expressing neurons were exposed to an impermeant acidic buffer to quench sypHy fluorescence on the presynaptic plasma membrane, since the altered surface fraction of SV cargo can be indicative of chronic dysfunction in either their clustering or retrieval (Gordon et al., 2011; Pan et al., 2015). Second, after recovery of the sypHy response to the acid buffer, neurons were challenged with two sequential AP trains (10 Hz, 30 s or 40 Hz 10 s) to reveal whether SV fusion or cargo retrieval was disrupted during periods of low and high activity. Finally, neurons were exposed to buffer containing NH_4_Cl, to unquench all sypHy within acidic compartments, to estimate the extent of SV fusion as a proportion of the total SV pool. In parallel, activity-dependent uptake of the large fluid phase marker tetramethylrhodamine (TMR)-dextran (40 kDa), was measured, since it exclusively reports ADBE due to size exclusion from SVs (Harper & Smillie, 2021).

### ADBE is depressed in both Nrxn1^+/−^ and Nlgn3^-/y^ neurons

We first tested our high content imaging protocol on two preclinical rodent models - *Nrxn1^+/−^* and *Nlgn3^-/y^*rats. These rat models replicate the genetic alterations in the human *NRXN1* or *NLGN3* genes that are primarily responsible for ASDs (Südhof, 2017; Uchigashima et al., 2021), due to haploinsufficiency of the *Nrxn1* gene, or deletion of the X-linked *Nlgn3* gene respectively. An additional reason for investigating these rat models is that an unbiased screen of molecules on bulk endosomes generated via ADBE revealed that synaptic adhesion molecules were disproportionately represented (Kokotos et al., 2018). Furthermore, the gene products of *Nrxn1* and *Nlgn3*, Neurexin-1 and Neuroligin-3, were present on bulk endosomes purified via two independent approaches (Kokotos et al., 2018). This suggests the potential for a direct mechanistic link between adhesion molecules, ASDs and ADBE. Therefore, we first determined whether these construct-valid rat models of ASDs displayed dysfunctional SV recycling and ADBE.

Neurexins are presynaptic cell adhesion molecules that stabilise synapses via trans-synaptic interactions with postsynaptic partners such as neuroligins (Südhof, 2017). Their conditional knockout has pleotropic effects on evoked neurotransmitter release, with the primary deficit being an inefficient coupling of voltage-gated calcium influx to SV fusion (Chen et al., 2017; Luo et al., 2020). Since ADBE is also triggered via activity-dependent calcium influx (Clayton et al., 2009; Morton et al., 2015), we first determined the impact of *Nrxn1* haploinsufficiency on ADBE and SV recycling in primary hippocampal cultures from the *Nrxn1^+/−^* rat (Achterberg et al., 2024; Kight et al., 2021). When the high content sypHy assay was performed on *Nrxn1^+/−^* neurons and wild-type littermate controls, no significant difference in the extent of the evoked sypHy response was observed to either AP train (10 Hz or 40 Hz, Figure 2A,C,D,E). This suggests that SV fusion was not significantly impacted by the absence of a single *Nrxn1* allele. Furthermore, the kinetics and extent of SV cargo retrieval in *Nrxn1^+/−^*neurons were not significantly impacted when compared to wild-type neurons (Figure 2F,G). This absence of effect was corroborated by the comparable extent of sypHy surface fraction between wild-type and *Nrxn1^+/−^* neurons (Figure 2B). However, when ADBE was assessed, a significant reduction in the number of nerve terminals displaying activity-dependent uptake of TMR-dextran was observed in *Nrxn1^+/−^*neurons when compared to wild-type littermate controls (Figure 2H,I). This decrease in TMR-dextran puncta number was not due to a reduction in nerve terminals, since staining with the SV marker SV2A revealed no change in this parameter when *Nrxn1^+/−^* and wild-type neurons were compared (Figure 2J,K). Therefore *Nrxn1^+/−^* neurons display a selective deficit in ADBE, but not SV fusion or cargo retrieval.

**Figure 2.**
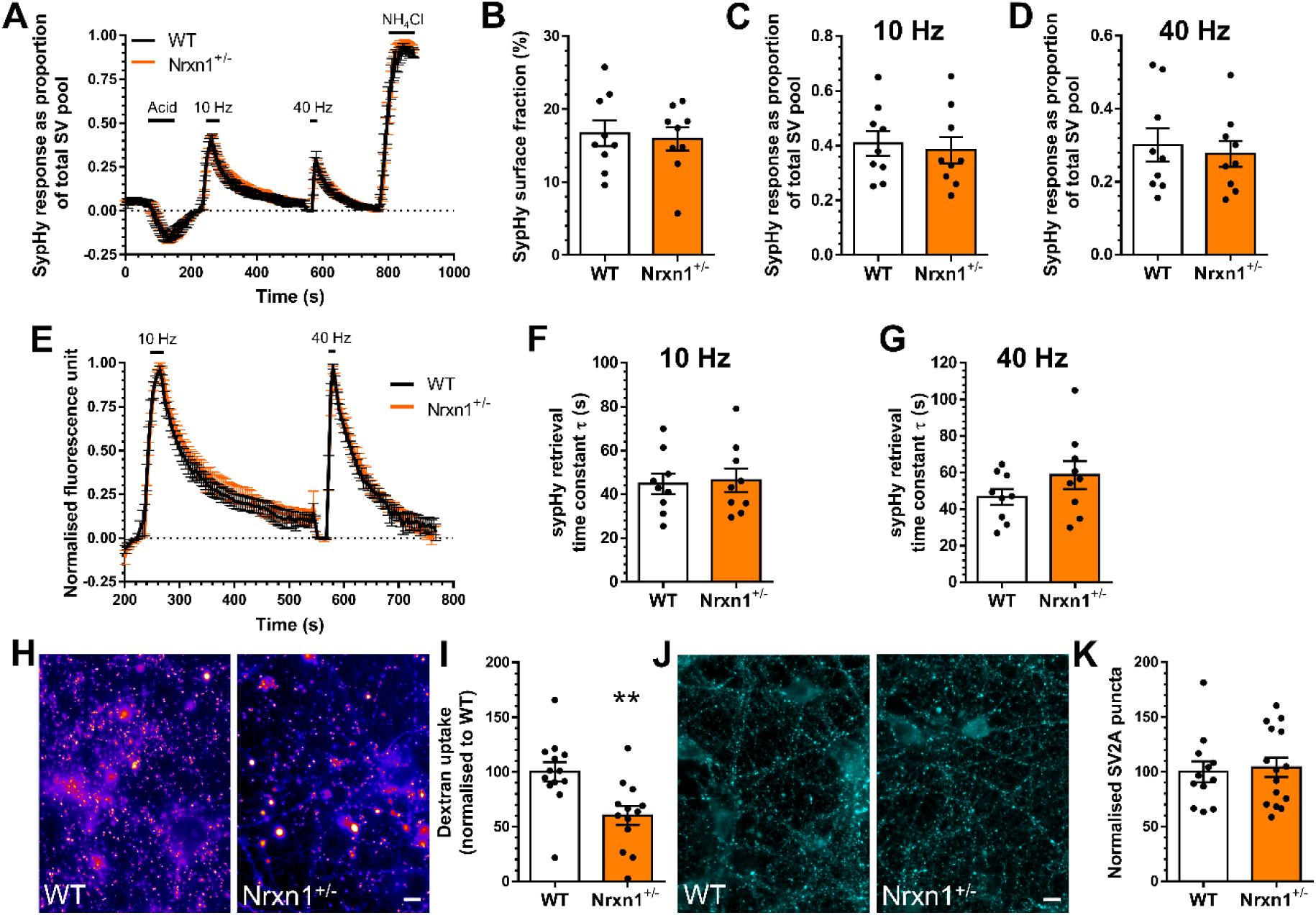
*Nrxn1^+/−^*neurons display depressed ADBE. Hippocampal neurons derived from either wild-type (WT) or *Nrxn1^+/−^* rat embryos were transfected with synaptophysin-pHluorin (sypHy) after 7 days *in vitro* (DIV) and imaged DIV 13-15. A, E) Mean sypHy fluorescence traces of WT (black) and *Nrxn1^+/−^* (orange) hippocampal neurons normalised to either the total SV pool as revealed by NH_4_Cl (A) or peak fluorescence during electrical stimulation (E) ± SEM. B) Mean sypHy surface fraction presented as a percentage of the total SV pool ± SEM. C,D) Mean peak sypHy response in response to either 10 Hz (C) or 40 Hz (D) action potential trains ± SEM. F,G) Mean sypHy retrieval time constants (τ) in response to either 10 Hz (F) or 40 Hz (G) action potential trains ± SEM. For A-G WT n= 9 coverslips, Het n = 9 from 3 independent cultures. H-I) Primary hippocampal cultures derived either wild-type (WT) or *Nrxn1^+/−^* rat embryos were challenged with a train of action potentials (40 Hz, 10 s) in the presence of tetramethylrhodamine (TMR)-dextran (50 μM). TMR-dextran was immediately washed away and the number of dextran puncta were counted. H) Representative images of dextran uptake in WT and *Nrxn1^+/−^* cultures. Scale bar = 50µm. I) Mean number of TMR- dextran puncta per field of view normalised to WT ± SEM (n= 13 coverslips from 3 independent cultures for WT and *Nrxn1^+/−^*). J) Primary hippocampal cultures derived from either WT or *Nrxn1^+/−^*rat embryos were fixed at DIV13-15 and stained for the presence of SV2A. K) Mean number of SV2A puncta per field of view normalise to WT ± SEM (WT n= 12 coverslips, *Nrxn1^+/−^* n = 15 coverslips from 3 independent cultures). In all cases an unpaired two-sided students t test was performed, ** p=0.0037, unpaired t test.

Neurexin-1 forms complexes with postsynaptic adhesion molecules to drive synaptogenesis and synaptic stability (Craig & Kang, 2007). Key proteins in this context are the neuroligin family, of which neuroligin-3 is an important member. Neuroligin-3 is located at both excitatory and inhibitory synapses and performs essential roles in synaptic development, function and maintenance (Varoqueaux et al., 2006). As stated above, *NLGN3* gene mutations are associated with ASDs, with the majority of pathogenic mutations resulting in a loss of neuroligin-3 function (Uchigashima et al., 2021). Therefore, we next determined whether ADBE was depressed in hippocampal neurons derived from a recently generated model of neuroligin-3 dysfunction, the *Nlgn3^-/y^*rat (Anstey et al., 2022). We first addressed whether *Nlgn3^-/y^*neurons displayed altered SV fusion or cargo retrieval using our high content sypHy screen. *Nlgn3^-/y^* neurons displayed no defect in SV fusion in response to either stimulation protocol, when compared to wild-type littermate controls (Figure 3A,C,D,E). Furthermore, there was no difference between *Nlgn3^-/y^* and wild-type neurons in terms of either the extent or kinetics of sypHy retrieval (Figure 3F,G). Finally, there was no significant change in the surface fraction of sypHy between the two genotypes (Figure 3B), suggesting both SV cargo retrieval and clustering are unaffected by the absence of neuroligin-3. In contrast, the number of nerve terminals displaying an activity-dependent accumulation of TMR-dextran was reduced in *Nlgn3^-/y^* neurons when compared to wild-type controls (Figure 3H,I). Since the number of nerve terminals was unchanged in *Nlgn3^-/y^* cultures (revealed by SV2A immunostaining, Figure 3J,K), this meant that, similar to *Nrxn1^+/−^* neurons, *Nlgn3^-/y^*neurons display reduced ADBE.

**Figure 3.**
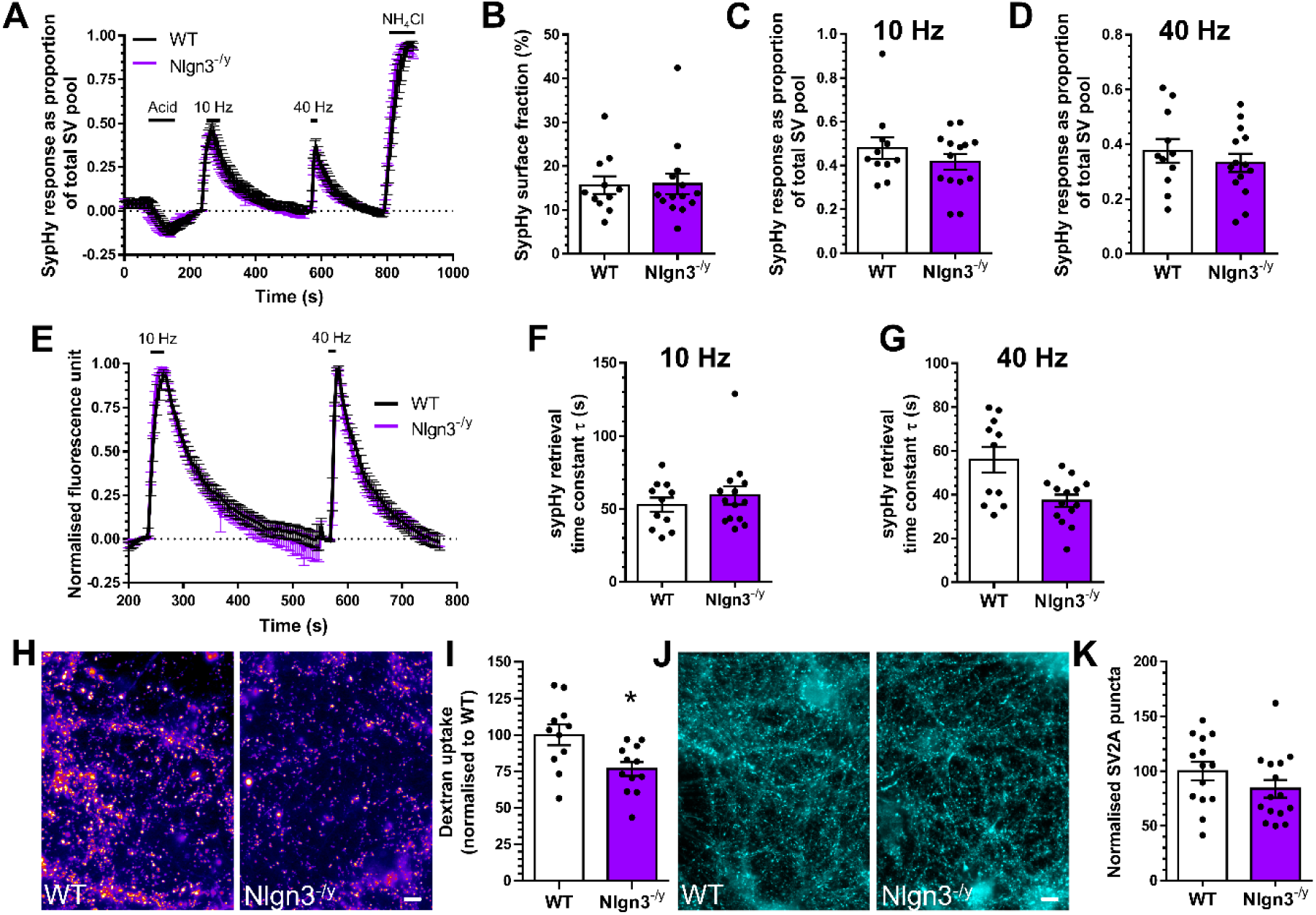
*Nlgn3^-/y^*neurons display depressed ADBE. Hippocampal neurons derived from either wild-type (WT) or *Nlgn3^-/y^* rat embryos were transfected with synaptophysin-pHluorin (sypHy) after 7 days *in vitro* (DIV) and imaged DIV 13-15. A, E) Mean sypHy fluorescence traces of WT (black) and *Nlgn3^-/y^* (purple) hippocampal neurons normalised to either the total SV pool as revealed by NH_4_Cl (A) or peak fluorescence during electrical stimulation (E) ± SEM. B) Mean sypHy surface fraction presented as a percentage of the total SV pool ± SEM. C,D) Mean peak sypHy response in response to either 10 Hz (C) or 40 Hz (D) action potential trains ± SEM. F,G) Mean sypHy retrieval time constants (τ) in response to either 10 Hz (F) or 40 Hz (G) action potential trains ± SEM. For A-G WT n= 11 coverslips, *Nlgn3^-/y^* n = 14 from 3 independent cultures. H-I) Primary hippocampal cultures derived either wild-type (WT) or *Nlgn3^-/y^*rat embryos were challenged with a train of action potentials (40 Hz, 10 s) in the presence of tetramethylrhodamine (TMR)-dextran (50 μM). TMR-dextran was immediately washed away and the number of dextran puncta were counted. H) Representative images of dextran uptake in WT and *Nlgn3^-/y^* cultures. Scale bar = 50µm. I) Mean number of TMR- dextran puncta per field of view normalised to WT ± SEM (WT n= 11 coverslips, *Nlgn3^-/y^* n = 12 from 3 independent cultures). J) Primary hippocampal cultures derived from either WT or *Nlgn3^-/y^* rat embryos were fixed at DIV13-15 and stained for the presence of SV2A. K) Mean number of SV2A puncta per field of view normalise to WT ± SEM (WT n= 14 coverslips, *Nlgn3^-/y^* n = 15 coverslips from 3 independent cultures). In all cases an unpaired two-sided students t test was performed, except B,F and G, * p=0.0112, unpaired t test.

### Loss of SynGAP results in depressed ADBE

Neurexin-1 is a presynaptic protein, whereas neuroligin-3 forms essential complexes that modify and maintain presynaptic function (Craig & Kang, 2007). Therefore, the observed depression of ADBE may result from a direct mechanistic involvement of these molecules in this endocytosis mode. To test the hypothesis that depression in ADBE was a consequence of the brain microenvironment in ASDs, and not to loss of a key regulatory molecule, we next examined ASD models where the affected gene product is expressed exclusively at the postsynapse. The models chosen for these experiments were the *Syngap*^+/−^ rat and the *Syngap*^+/Δ-GAP^ rat (Buller-Peralta et al., 2022; Katsanevaki et al., 2024; Mastro et al., 2020). SynGAP performs key roles at the postsynapse, principally in the control of AMPA receptor trafficking and synaptic plasticity (Gamache et al., 2020). Furthermore, *SYNGAP* haploinsufficiency is a highly prevalent cause of ASD, resulting from loss of function mutations in the *SYNGAP* gene (Mignot et al., 2016; Vlaskamp et al., 2019). The *Syngap*^+/−^ rat model was generated by introducing a frameshift mutation in the *Syngap* gene, resulting in nonsense mediated decay of mRNA encoding the mutant allele (Masto et al 2020). In contrast, the *Syngap*^+/Δ-GAP^ rat has a deletion in the exons encoding the calcium/lipid binding C2, and GTPase activating GAP domain (Buller-Peralta et al., 2022; Katsanevaki et al., 2024).

As stated above, the exclusive postsynaptic localisation of SynGAP provides an excellent opportunity to test the hypothesis that depression of ADBE is a consequence of the ASD brain microenvironment. We first determined the ability of *Syngap*^+/−^ hippocampal neurons to perform efficient SV fusion and cargo trafficking using our sypHy assay. In this instance, we also examined the performance of *Syngap*^-/−^ neurons in this assay, to determine if the complete absence of the protein exacerbated any potential phenotype. When SV fusion was assessed, neither *Syngap*^+/−^ nor *Syngap*^-/−^ neurons displayed any deficit when compared to wild-type littermate controls during either the low or high frequency stimulus train (Figure 4A,C,D,E). Furthermore, neither genotype displayed dysfunctional SV cargo retrieval in response to these trains, or an alteration in the surface distribution of sypHy (Figure 4B,F,G). In contrast, *Syngap*^+/−^ neurons displayed a reduction the number of activity-dependent TMR-dextran puncta, a decrease which became significant in *Syngap*^-/−^ neurons when compared to wild-type controls (Figure 4H,I). This was not a result of a decrease in nerve terminals in these primary cultures, since the number of SV2A puncta was unchanged across all genotypes (Figure 4J,K). Therefore, a depression in ADBE is still observed even in neurons where the affected gene product is exclusively postsynaptic.

**Figure 4.**
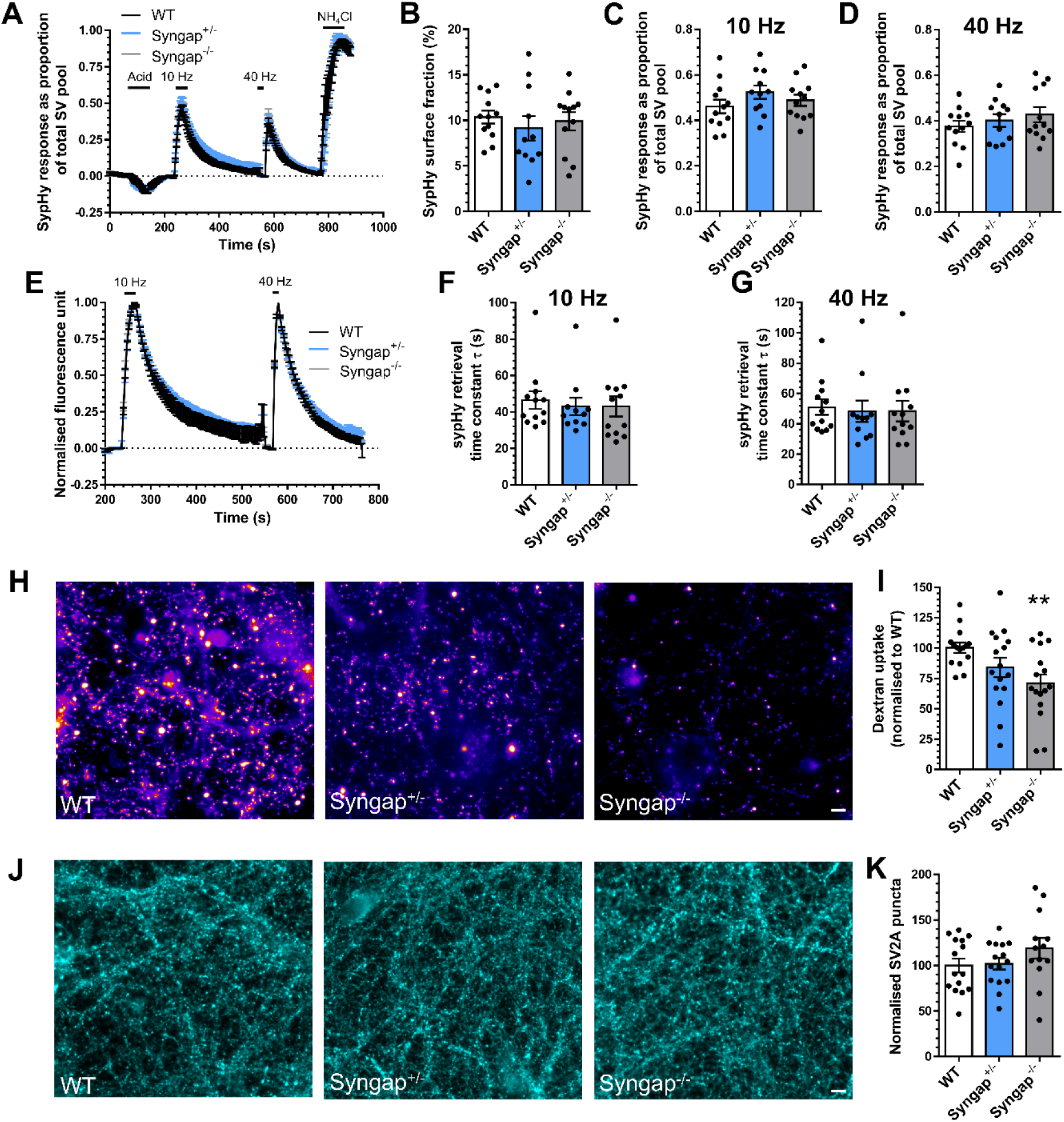
*Syngap*^-/−^ neurons display depressed ADBE. Hippocampal neurons derived from either wild-type (WT), *Syngap*^+/−^ or *Syngap*^-/−^ rat embryos were transfected with synaptophysin-pHluorin (sypHy) after 7 days *in vitro* (DIV) and imaged DIV 13-15. A, E) Mean sypHy fluorescence traces of WT (black), *Syngap*^+/−^ (blue) or *Syngap*^-/−^ (grey) hippocampal neurons normalised to either the total SV pool as revealed by NH_4_Cl (A) or peak fluorescence during electrical stimulation (E) ± SEM. B) Mean sypHy surface fraction presented as a percentage of the total SV pool ± SEM. C,D) Mean peak sypHy response in response to either 10 Hz (C) or 40 Hz (D) action potential trains ± SEM. F,G) Mean sypHy retrieval time constants (τ) in response to either 10 Hz (F) or 40 Hz (G) action potential trains ± SEM. For A-G, WT n= 12 coverslips, *Syngap*^+/−^ n = 11 and *Syngap*^-/−^ n = 12 from 3 independent cultures. H-I) Primary hippocampal cultures derived either wild-type (WT), *Syngap*^+/−^ or *Syngap*^-/−^ rat embryos were challenged with a train of action potentials (40 Hz, 10 s) in the presence of tetramethylrhodamine (TMR)-dextran (50 μM). TMR-dextran was immediately washed away and the number of dextran puncta were counted. H) Representative images of dextran uptake in WT, *Syngap*^+/−^ or *Syngap*^-/−^ cultures. Scale bar = 50µm. I) Mean number of TMR-dextran puncta per field of view normalised to WT ± SEM (WT n= 15 coverslips, *Syngap*^+/−^ and *Syngap*^-/−^ n = 16 from 3 independent cultures). J) Primary hippocampal cultures derived from either WT, *Syngap*^+/−^ or *Syngap*^-/−^ rat embryos were fixed at DIV13-15 and stained for the presence of SV2A. K) Mean number of SV2A puncta per field of view normalise to WT ± SEM (WT and *Syngap*^+/−^ n = 15 coverslips, *Syngap*^-/−^ n = 13 from 3 independent cultures). In all cases a one-way ANOVA was performed, ** p=0.01.

SynGAP functions as a GAP that negatively regulates Ras and Rap GTPases to control both F-actin dynamics (RasGAP) and p38 MAPK (RapGAP) activity (Chen et al., 1998; Krapivinsky et al., 2004; Pena et al., 2008). However, SynGAP has other functions outside of its GAP activity. Its C-terminus alters synaptic strength via PDZ interactions with PSD-95, which in turn control AMPA receptor recruitment and synaptogenesis (Kim et al., 1998; Opazo et al., 2012; Walkup et al., 2016). While the role of the GAP activity in brain function remains poorly understood, recent work has suggested that this catalytic domain may not be obligatory for synaptic plasticity (Araki et al., 2024). To delineate these more structural roles from its GAP activity, we next exploited the *Syngap*^+/Δ-GAP^ rat model (Buller-Peralta et al., 2022; Katsanevaki et al., 2024), which shares many behavioural traits with the *Syngap*^+/−^ rat. Primary hippocampal cultures were prepared from both *Syngap*^+/Δ-GAP^ and *Syngap^Δ-GAP^*^/Δ-GAP^ rats in addition to wild-type littermate controls. When the three genotypes were assessed for SV fusion phenotypes using the sypHy assay, there was no significant difference for either stimulus train (Figure 5A,C,D,E). There was also no significant effect of genotype on SV cargo retrieval after either stimulus and no impact on the surface fraction of sypHy (Figure 5B,F,G). When the number of activity-dependent TMR-dextran puncta were determined, *Syngap*^+/Δ-GAP^ neurons displayed a reduction, which again became significant when *Syngap^Δ-GAP^*^/Δ-GAP^ neurons were examined (Figure 5H,I). Furthermore, the number of nerve terminals in culture, identified via SV2A staining, was unchanged across all genotypes (Figure 5J,K). This result therefore confirms that loss of postsynaptic SynGAP function in two independent model systems results in depression of ADBE.

**Figure 5.**
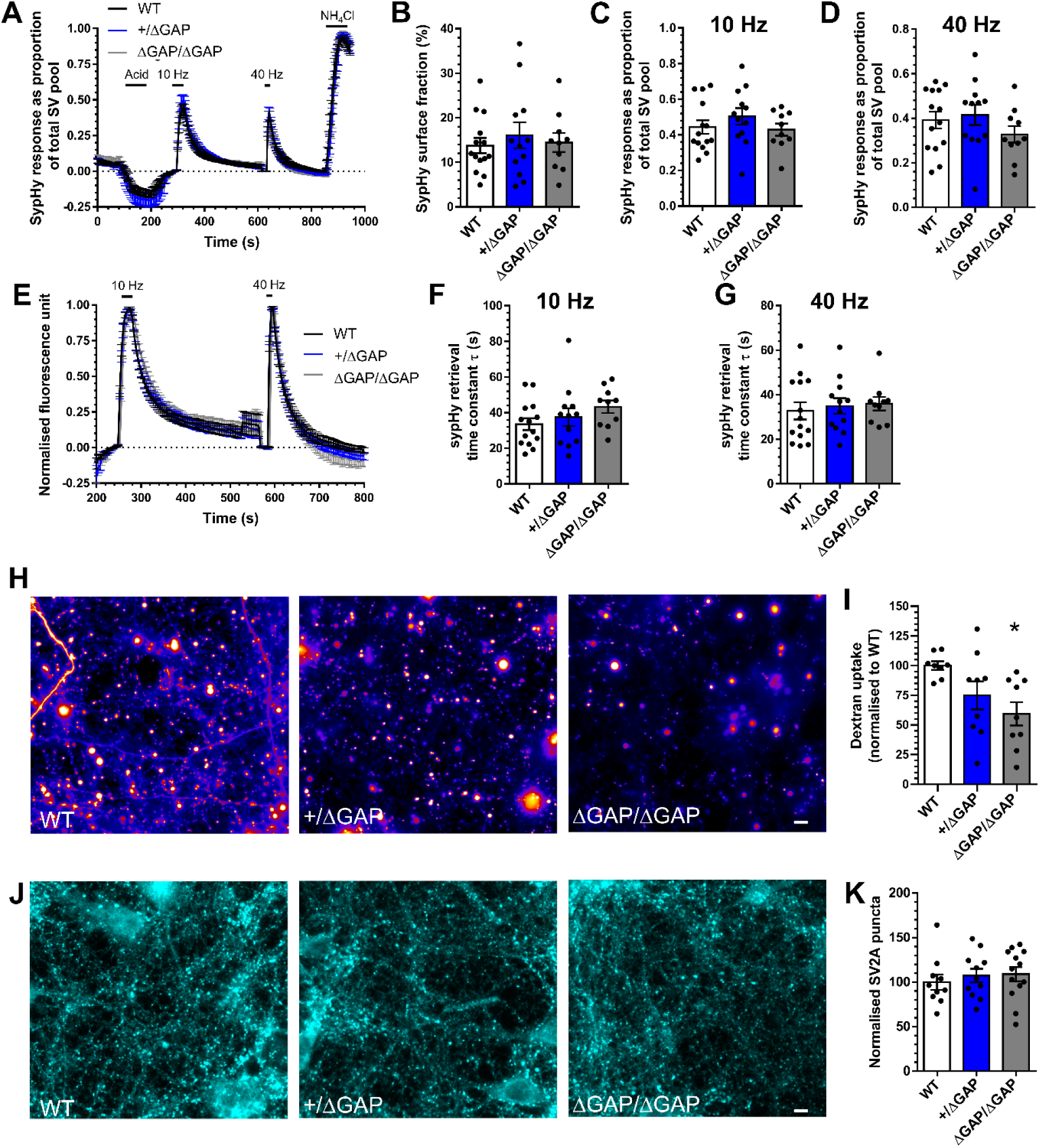
*Syngap^Δ-GAP^*^/Δ-GAP^ neurons display depressed ADBE. Hippocampal neurons derived from either wild-type (WT), *Syngap*^+/Δ-GAP^ or *Syngap^Δ-GAP^*^/Δ-GAP^ rat embryos were transfected with synaptophysin-pHluorin (sypHy) after 7 days *in vitro* (DIV) and imaged DIV 13-15. A, E) Mean sypHy fluorescence traces of WT (black), *Syngap*^+/Δ-GAP^ (blue) or *Syngap^Δ- GAP^*^/Δ-GAP^ (grey) hippocampal neurons normalised to either the total SV pool as revealed by NH_4_Cl (A) or peak fluorescence during electrical stimulation (E) ± SEM. B) Mean sypHy surface fraction presented as a percentage of the total SV pool ± SEM. C,D) Mean peak sypHy response in response to either 10 Hz (C) or 40 Hz (D) action potential trains ± SEM. F,G) Mean sypHy retrieval time constants (τ) in response to either 10 Hz (F) or 40 Hz (G) action potential trains ± SEM. For A-G, WT n= 14 coverslips, *Syngap*^+/Δ-GAP^ n = 12 and *Syngap^Δ-GAP^*^/Δ-GAP^ n = 10 from 3 independent cultures. H-I) Primary hippocampal cultures derived either wild-type (WT), *Syngap*^+/Δ-GAP^ or *Syngap^Δ-GAP^*^/Δ-GAP^ rat embryos were challenged with a train of action potentials (40 Hz, 10 s) in the presence of tetramethylrhodamine (TMR)-dextran (50 μM). TMR-dextran was immediately washed away and the number of dextran puncta were counted. H) Representative images of dextran uptake in WT, *Syngap*^+/Δ-GAP^ or *Syngap^Δ-GAP^*^/Δ-GAP^ cultures. Scale bar = 50µm. I) Mean number of TMR-dextran puncta per field of view normalised to WT ± SEM (WT n= 8 coverslips, *Syngap*^+/Δ-GAP^ and *Syngap^Δ-GAP^*^/Δ-GAP^ n = 9 from 3 independent cultures). J) Primary hippocampal cultures derived from either WT, *Syngap*^+/Δ-GAP^ or *Syngap^Δ-GAP^*^/Δ-GAP^ rat embryos were fixed at DIV13-15 and stained for the presence of SV2A. K) Mean number of SV2A puncta per field of view normalise to WT ± SEM (WT n = 10 coverslips, *Syngap*^+/Δ-GAP^ n = 11 and *Syngap^Δ-GAP^*^/Δ-GAP^ n = 13 from 3 independent cultures). In all cases a one-way ANOVA was performed, * p=0.017.

### Pten^+/−^ neurons display depressed ADBE

The demonstration of depression of ADBE in SynGAP models, which have exclusively postsynaptic deficits, provides strong support for the hypothesis that a reduction in ADBE is a convergent consequence of ASD. As a final proof, we exploited a different rat ASD model that has no discernible presynaptic locus of dysfunction, the *Pten*^+/−^ rat (Rowley et al., 2019). PTEN (phosphatase and tensin homologue deleted on chromosome 10) is a tumour suppressor, that negatively regulates the AKT / mTOR signalling pathway (Waite & Eng, 2002). Haploinsufficiency resulting from loss of function mutations in the *PTEN* gene result in PTEN hamartoma tumor syndrome (PHTS, (Busch et al., 2019)), with common links to ASDs, macrocephaly, epilepsy and neurodevelopmental impairment (Conti et al., 2012; Klein et al., 2013; Satterstrom et al., 2020).

Primary hippocampal cultures from either *Pten*^+/−^ rat embryos or wild-type littermate controls were entered into the sypHy high content assay. Consistent with other models of ASD, *Pten*^+/−^ neurons displayed no defect in SV fusion, cargo retrieval or surface stranding of sypHy (Figure 6A-G). Intriguingly, *Pten*^+/−^ neurons did display a reduction in the number of activity-dependent TMR-dextran puncta when compared to littermate controls (Figure 6H,I). Therefore, in an ASD model system with no overt presynaptic dysfunction, depression of ADBE still occurs.

**Figure 6.**
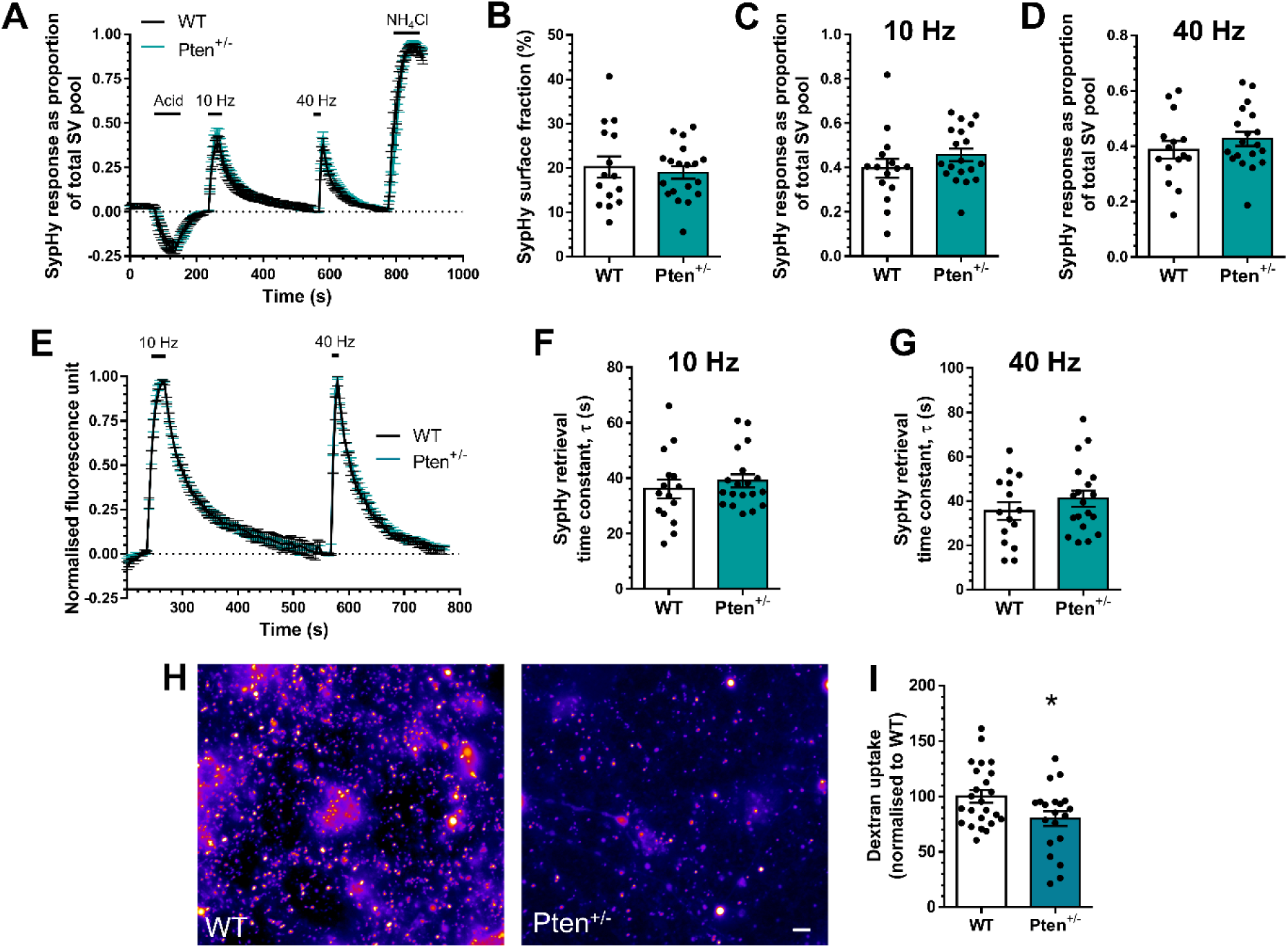
*Pten^+/−^*neurons display depressed ADBE. Hippocampal neurons derived from either wild-type (WT) or *Pten^+/−^* rat embryos were transfected with synaptophysin-pHluorin (sypHy) after 7 days *in vitro* (DIV) and imaged DIV 13-15. A, E) Mean sypHy fluorescence traces of WT (black) and *Pten^+/−^*(turquoise) hippocampal neurons normalised to either the total SV pool as revealed by NH_4_Cl (A) or peak fluorescence during electrical stimulation (E) ± SEM. B) Mean sypHy surface fraction presented as a percentage of the total SV pool ± SEM. C,D) Mean peak sypHy response in response to either 10 Hz (C) or 40 Hz (D) action potential trains ± SEM. F,G) Mean sypHy retrieval time constants (τ) in response to either 10 Hz (F) or 40 Hz (G) action potential trains ± SEM. For A-G WT n= 15 coverslips, *Pten^+/−^* n = 19 from 3 independent cultures. H-I) Primary hippocampal cultures derived either wild-type (WT) or *Pten^+/−^*rat embryos were challenged with a train of action potentials (40 Hz, 10 s) in the presence of tetramethylrhodamine (TMR)-dextran (50 μM). TMR-dextran was immediately washed away and the number of dextran puncta were counted. H) Representative images of dextran uptake in WT and *Pten^+/−^* cultures. Scale bar = 50µm. I) Mean number of TMR- dextran puncta per field of view normalised to WT ± SEM (WT n= 24 coverslips, *Pten^+/−^* n = 20 from 3 independent cultures).

## Discussion

The complex aetiology of ASDs complicate the investigation of their causal mechanisms. Because of this, the study of monogenic ASDs, where the genetic locus is known, has provided a series of key insights into potential convergent signalling pathways (Yuen et al., 2017). In this study we have revealed that neurons derived from a range of rodent models of monogenic ASDs display a depression of ADBE, a presynaptic endocytosis mode that is dominant during periods of high neuronal activity. This depression was observed regardless of whether the mutated gene was presynaptic, postsynaptic or neither, suggesting it is a convergent strategy to ameliorate disrupted brain function.

We initially revealed depression in ADBE in *Fmr1^-/y^*neurons, a model for fragile X syndrome (Bonnycastle et al., 2022). Similar to this study, SV fusion and cargo trafficking were unaffected. A BK channel agonist corrected the depression of ADBE in these neurons, suggesting the alteration could be a result of the direct regulation of BK channels by the *Fmr1* gene product, FMRP (Deng & Klyachko, 2016; Deng et al., 2013; Myrick et al., 2015). However, a BK channel antagonist did not recapitulate the depression in wild-type neurons, suggesting that FMRP had no direct mechanistic role in ADBE (Bonnycastle et al., 2022). In this work, we determined whether ADBE was perturbed in other ASD models that did not display overt defects in presynaptic endocytosis. The large fluid phase marker TMR-dextran was used for these studies, since it reports ADBE due to size exclusion from single SVs (Clayton et al., 2008). TMR-dextran is an excellent reporter of ADBE, since interventions that block this endocytosis mode selectively reduce TMR-dextran uptake (Blumrich et al., 2023; Clayton et al., 2009; Clayton et al., 2010; Morton et al., 2015; Smillie et al., 2013).

Specific cell adhesion molecules such as CHL1 and N-cadherins have proposed roles at the presynapse in both activity-dependent SV retrieval and ADBE (Dagar et al., 2021; Leshchyns’ka et al., 2006; van Stegen et al., 2017; Vitureira et al., 2011). In fact, cell adhesion molecules including both neurexin-1 and neuroligin-3 are greatly over-represented on ADBE-generated bulk endosomes (Kokotos et al., 2018). This suggests that they might be required for optimal ADBE. In support, neurexin-1 controls localised calcium channel coupling to SV fusion at the active zone (Chen et al., 2017; Luo et al., 2020; Südhof, 2017) and both neurexin-1 and neuroligin-3 are synaptogenic (Craig & Kang, 2007). However, a direct role via these presynaptic functions is unlikely. This is because ADBE is triggered via delocalised calcium increases, rather than localised calcium influx at the active zone (Morton et al., 2015) and there was no significant change in synapse number in either *Nrxn1^+/−^* or *Nlgn3^-/y^* cultures, suggesting synaptogenesis was not modulating this effect (this study).

Two independent models of *SYNGAP* haploinsufficiency disorder displayed depression of ADBE. The exclusive postsynaptic location of SynGAP provides strong support for the depression of ADBE being a compensatory adjustment in ASDs. The expression of specific isoforms of SynGAP are driven via neuronal activity and have downstream effects on mEPSC frequency (McMahon et al., 2012), suggesting a potential presynaptic role. However, it is more likely that these effects are mediated via the nano-organisation of AMPA receptors at the postsynapse. The observation that a depression of ADBE was observed in *Syngap*^+/Δ-GAP^ neurons suggests that this depression resulted from a loss of enzyme activity, rather than interactions with postsynaptic partners such as PSD-95, LRRTMs and neuroligins (Kim et al., 1998; Opazo et al., 2012; Walkup et al., 2016). In this context key enzymatic roles for SynGAP could include either its RasGAP, which regulates F-actin and spine dynamics (Bär et al., 2016) or Rap-GAP activity which controls the p38 MAPK signalling pathway ultimately regulating postsynaptic AMPA receptor trafficking (Krapivinsky et al., 2004; Rumbaugh et al., 2006; Zhu et al., 2002). The obligatory role of the GAP domain in SYNGAP function has recently been questioned (Araki et al., 2024), however the GAP rat displays a series of circuit and behavioural phenotypes that suggest its enzymatic role of required for optimal function (Katsanevaki et al., 2024). Regardless, the depression of ADBE in two independent models where the gene product is expressed exclusively at the postsynapse provides strong support for this depression to be a homeostatic adaptation to circuit hyperexcitability.

The depression of ADBE in a novel *Pten*^+/−^ model could potentially be explained by the fact that PTEN is expressed at growth cones during axonal navigation and synaptogenesis. However, its expression is restricted to the postsynapse in mature neurons (Kreis et al., 2014). The synaptic locus of dysfunction in neurons lacking PTEN appears to be due to dysregulation of the AKT/mTOR signalling pathway, specifically mTORC1/RAPTOR (Tariq et al., 2022). Deficiency in PTEN results in neuronal hypertrophy and hyperexcitability (Santos et al., 2017; Skelton et al., 2019; Southwell et al., 2020; Williams et al., 2015), which is corrected in multiple models via inhibition of AKT/mTOR signalling via rapamycin (Getz et al., 2016; Tariq et al., 2022; Weston et al., 2012). PTEN deficient neurons display enhanced excitatory neurotransmission, which appears to be due to enhanced postsynaptic function. Intriguingly, an increase in the size of the RRP and mEPSC frequency was also observed, however this was most likely due to an increase in the number of available synapses in these neurons (Tariq et al., 2022; Weston et al., 2012; Williams et al., 2015). In agreement with this hypothesis, no change in evoked SV fusion events were detected in this study.

The central question arising from this study is, what the molecular mechanism responsible for the depression of ADBE in ASD model neurons? ADBE is the dominant endocytosis mode during high activity and appear to have a similar molecular mechanism to ultrafast endocytosis, which is triggered by sparse stimulation (Clayton et al., 2009; Clayton et al., 2008; Imoto et al., 2022; Watanabe et al., 2018; Watanabe et al., 2013). Since both pathways occur with a timescale that is an order of magnitude faster than SV cargo clustering and retrieval via CME, an emerging view is that CME initiates on the presynaptic plasma membrane but completes on endosomes formed via either UFE or ADBE (Cousin, 2017; Kononenko et al., 2014; Watanabe et al., 2014). Since SV fusion and cargo retrieval are not impacted in ASD models, it suggests that the depression is intrinsic to ADBE or mechanisms that control it. One compelling explanation is that the observed depression is a compensatory mechanism to limit the intrinsic circuit hyperexcitability observed in many ASD models including many of the models used in this study (Avazzadeh et al., 2021; Booker et al., 2019; Clement et al., 2012; Das Sharma et al., 2020; Modi et al., 2019; Ozkan et al., 2014; Santos et al., 2017; Skelton et al., 2019; Southwell et al., 2020; Tariq et al., 2022; Williams et al., 2015) but see (Michaelson et al., 2018). It will be key to determine the impact of elevated excitability in the long-term on presynaptic events such as ADBE.

Out of all potential presynaptic endocytosis modes, a depression in ADBE is the most intriguing, since its depression would be manifested disproportionately across specific cell types. This is because it is triggered during periods of high activity (Clayton et al., 2008), meaning that GABAergic neurons and inhibitory circuits (which typically display higher firing rates (Bartos et al., 2007), may be disproportionately impacted. In agreement, reduced ADBE results in an inability to sustain neurotransmission during high activity (Blumrich et al., 2023; Bonnycastle et al., 2022; Nicholson-Fish et al., 2015). Therefore, a depression of ADBE may not be a homogenous adaptation across the brain, but rather a circuit-specific correction in neuronal function. It will be essential in future work to determine how, 1) ADBE is impacted in intact circuits in ASD model system and 2) how circuit-specific modulation of this activity-dependent endocytosis mode may sculpt ASD-like behaviours.

## Materials and Methods

### Materials

Unless otherwise specified, all cell culture reagents were obtained from Invitrogen (Paisley, UK). Foetal bovine serum was from Biosera (Nuaille, France). Papain was obtained from Worthington Biochemical (Lakewood, NJ, USA). All other reagents were obtained from Sigma-Aldrich (Poole, UK) unless specified. Rabbit anti-SV2A was obtained from Abcam (Cambridge, UK; ab32942 RRID: AB_778192). Anti-rabbit Alexa Fluor 488 (A11008 RRID: AB_143165) was obtained from Invitrogen (Paisley, UK). Synaptophysin-pHluorin was a gift from Prof. L. Lagnado (University of Sussex, UK).

### Rat models

Procedures were performed in accordance with the UK Animal (Scientific Procedures) Act 1986, under Project and Personal Licence authority and were approved by the Animal Welfare and Ethical Review Body at the University of Edinburgh (Home Office project licence – 7008878). Similarly, procedures were conducted in accordance with protocols approved by the Institutional Animal Ethics Committee of Institute for Stem Cell Science and Regenerative Medicine, Bangalore. All animals were killed by Schedule 1 procedures in accordance with UK Home Office Guidelines; adults were killed by exposure to CO_2_ followed by decapitation, whereas embryos were killed by decapitation followed by destruction of the brain. In Edinburgh, rats were housed on a 12/12 h light/dark cycle with a 21 ± 2 °C room temperature and food/water ad libitum. In Bangalore, rats were maintained on a 14h light/10h dark cycle with *ad-libitum* access to diet and water.

All transgenic rats in this study were generated by Horizon Discovery (now Envigo). Sprague–Dawley *Nlgn3*^−*/y*^ transgenic rats were created as previously described (Anstey et al., 2022). Long Evans-*SG^em1/PWC^*, referred to in the manuscript as *Syngap^+/−^* rats, were created as described (Mastro et al., 2020). Colony founders of Long Evans-*SG^em2/PWC^*, referred to as *Syngap^+/Δ-GAP^*, were produced by zinc finger nuclease-mediated deletion of the GAP domain of *Syngap* as described (Buller-Peralta et al., 2022; Katsanevaki et al., 2024). Sprague Dawley Nrxn1tm1sage rats, referred to as *Nrxn1^+/−^*rats (Achterberg et al., 2024; Kight et al., 2021) were purchased from Horizon Discovery. Sprague Dawley *Pten^+/−^* rats were purchased from Horizon Discovery and were generated as described previously (Rowley et al., 2019).

### Hippocampal cultures

Hippocampi from each embryo (e18.5-e19.5) were processed separately to avoid contamination across genotypes. For rats with an X chromosome mutation (*Nlgn3*), male embryos were taken for hippocampal dissection. For all other rat lines embryos of both sexes were used.

Dissociated primary hippocampal cultures were prepared from embryos as previously described (Bonnycastle et al., 2022). Briefly, isolated hippocampi were digested in a 10 U/mL papain solution (Worthington Biochemical, LK003178) at 37°C for 20 min. The papain was then neutralised using DMEM F12 (ThermoFisher Scientific, 21331-020) supplemented with 10 % Foetal bovine serum (BioSera, S1810-500) and 1 % penicillin/streptomycin (ThermoFisher Scientific, 15140-122). Cells were triturated to form a single cell suspension and plated at 5 x 10^4^ cells (with the exception of single cell tetramethylrhodamine (TMR)-dextran uptake experiments, 2.5 x 10^4^ cells) per coverslip on laminin (10 µg/ mL; Sigma Aldrich, L2020) and poly-D-lysine (Sigma Aldrich, P7886) coated 25 mm glass coverslips (VWR International Ltd, Lutterworth, UK). Cultures were maintained in Neurobasal media (ThermoFisher Scientific, 21103-049) supplemented with 2 % B-27 (ThermoFisher Scientific, 17504-044), 0.5 mM L-glutamine (ThermoFisher Scientific, 25030- 024) and 1% penicillin/streptomycin. After 2-3 days *in vitro* (DIV), 1 µM of cytosine arabinofuranoside (Sigma Aldrich, C1768) was added to each well to inhibit glial proliferation. Hippocampal neurons were transfected with sypHy at *DIV* 7 using Lipofectamine 2000 (ThermoFisher Scientific, 11668027) prior to imaging at *DIV* 13-15.

### High content screening of SV recycling using sypHy

SypHy-transfected neurons were visualised at 500 nm band pass excitation with a 515 nm dichroic filter and a long-pass >520 nm emission filter on a Zeiss Axio Observer D1 inverted epifluorescence microscope (Cambridge, UK). Images were captured using an AxioCam 506 mono camera (Zeiss) with a Zeiss EC Plan Neofluar 40x/1.30 oil immersion objective. Image acquisition was performed using Zen Pro software (Zeiss). Hippocampal cultures were mounted in a Warner Instruments (Hamden, CT, USA) imaging chamber with embedded parallel platinum wires (RC-21BRFS) while undergoing constant perfusion with imaging buffer (119 mM NaCl, 2.5 mM KCl, 2 mM CaCl_2_, 2 mM MgCl_2_, 25 mM HEPES, 30 mM glucose at pH 7.4, supplemented with 10 μM 6-cyano-7-nitroquinoxaline-2,3-dione (Abcam, Cambridge, UK, ab120271) and 50 μM DL-2-Amino-5-phosphonopentanoic acid (Abcam, Cambridge, UK, ab120044). Images were acquired at 4 s intervals. After acquisition of a 1 min baseline, neurons were challenged with an impermeant acidic buffer (69 mM NaCl, 2.5 mM KCl, 2 mM CaCl2, 2 mM MgCl2, 25 mM MES, 30 mM glucose at pH 5.5) for 1 min. After returning to imaging buffer for 2 min, cultures were challenged with two field stimuli (delivered using a Digitimer LTD MultiStim system-D330 stimulator, current output 100 mA, current width 1 ms) separated by 5 min. Neurons were first stimulated at 10 Hz for 30 s (300 APs) then 40 Hz for 10 s (400 APs. Finally, after a 3 min recovery period, alkaline buffer (50 mM NH_4_Cl substituted for 50 mM NaCl in imaging buffer) was used to reveal the maximal pHluorin response.

Time traces were analysed using the FIJI distribution of Image J (National Institutes of Health). Images were aligned using the Rigid body model of the StackReg plugin (https://imagej.net/StackReg). Nerve terminal fluorescence was measured using the Time Series Analyser plugin (https://imagej.nih.gov/ij/plugins/time-series.html). Regions of interest (ROIs) 5 pixels in diameter were placed over nerve terminals that responded to the electrical stimulus. A response trace was calculated for each cell by averaging the individual traces from each selected ROI. For sypHy time traces, fluorescence decay time constants (tau, τ, s) were calculated by fitting a monoexponential decay curve to data from the time point after the end of electrical stimulation.

### TMR-dextran uptake

TMR-dextran (ThermoFisher Scientific, D1842) uptake was performed as described previously (Bonnycastle et al., 2022). Neurons were mounted on a Zeiss Axio Observer D1 microscope as described above before challenging with 400 action potentials (40 Hz) in the presence of 50 µM of TMR-dextran (40,000 MW) in imaging buffer. The TMR-dextran solution was immediately washed away after stimulation terminated, and images were acquired using 556/25 nm excitation and 630/98 nm emission bandpass filters (Zeiss) while undergoing constant perfusion. Per coverslip of cells, 3-6 different fields of view were imaged. The TMR-dextran puncta in each image were quantified using the Analyze Particles plugin of Image J (NIH, https://imagej.nih.gov/ij/developer/api/ij/plugin/filter/ParticleAnalyzer.html) to select and count particles of 0.23-0.91 µm^2^. For all experiments, for each condition, at least one unstimulated coverslip was imaged to correct for the background level of TMR-dextran uptake.

### SV2A immunofluorescence staining

Immunofluorescence staining was performed as previously described (Bonnycastle et al., 2022). Briefly, neurons were fixed with 4 % paraformaldehyde (Sigma Aldrich, 47608) in PBS for 15 min. Excess paraformaldehyde was quenched with 50 mM NH_4_Cl in PBS. Cells were then permeabilized in 1 % bovine serum albumin (BSA; Roche Diagnostics GmbH, Germany, 10735078001) in PBS-Triton 0.1 % solution for 5 min and blocked in 1 % BSA in PBS at room temperature for 1 h. After blocking, cells were incubated in rabbit anti-SV2A (1:200 dilution) for 1 h, after which the cultures were washed with PBS and incubated with fluorescently conjugated secondary antibodies (anti-rabbit Alexa Fluor 488; 1:1000 dilution) for 1 h. The coverslips were mounted on slides for imaging with FluorSave (Millipore, Darmstadt, Germany, 345789). SV2A puncta were visualised at 500 nm band pass excitation with a 515 nm dichroic filter and a long-pass >520 nm emission filter on a Zeiss Axio Observer D1 inverted epifluorescence microscope (Cambridge, UK). Images were captured using an AxioCam 506 mono camera (Zeiss) with a Zeiss EC Plan Neofluar 40x/1.30 oil immersion objective. SV2A puncta in each image were quantified using the Analyze Particles plugin of Image J to select and count particles of 0.23-3.18 µm^2^.

### Experimental design and statistical analysis

Microsoft Excel (Microsoft, Washington, USA) and Prism 6 software (GraphPad software Inc., San Diego USA) were used for data processing and analysis. The experimenter was blinded to genotype during data acquisition and analysis. For all figures, results are presented with error bars as ± SEM, and the n for each condition represents the number of coverslips imaged. For all assays, cells were obtained from at least three independent cultures. In sypHy assays, at least 10 ROIs were collected from each coverslip. The number of ROIs examined was comparable for all experiments. Normality was determined using a D’Agostino & Pearson omnibus normality test. For comparison between genotypes, an unpaired Student t test was performed where data followed a normal distribution, with a Mann-Whitney test performed for those that did not. For comparison between >2 conditions, a one-way ANOVA was performed. Results were corrected for multiple comparisons. Full statistical reporting is provided in Table 1.

**Table 1.**
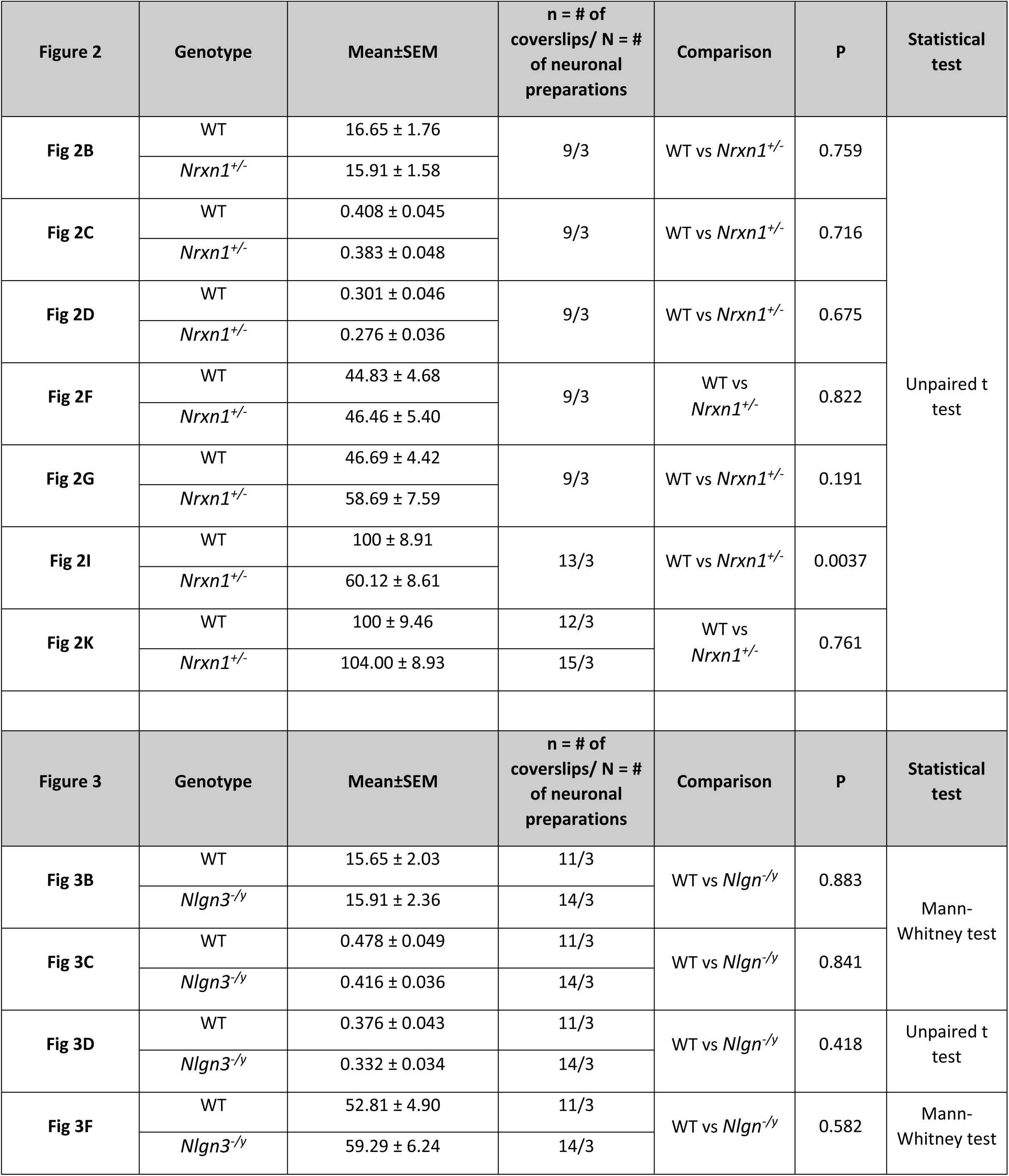
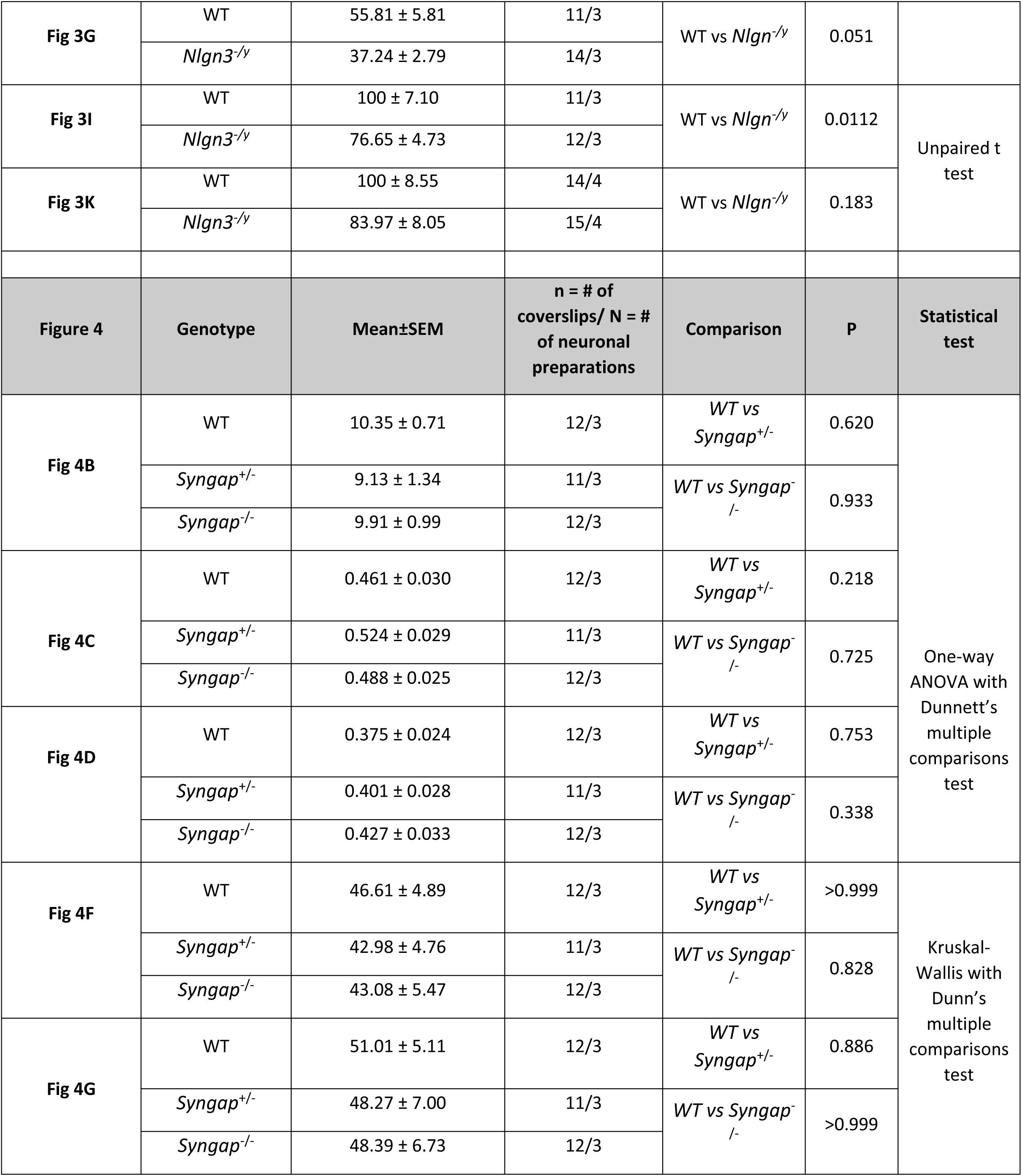
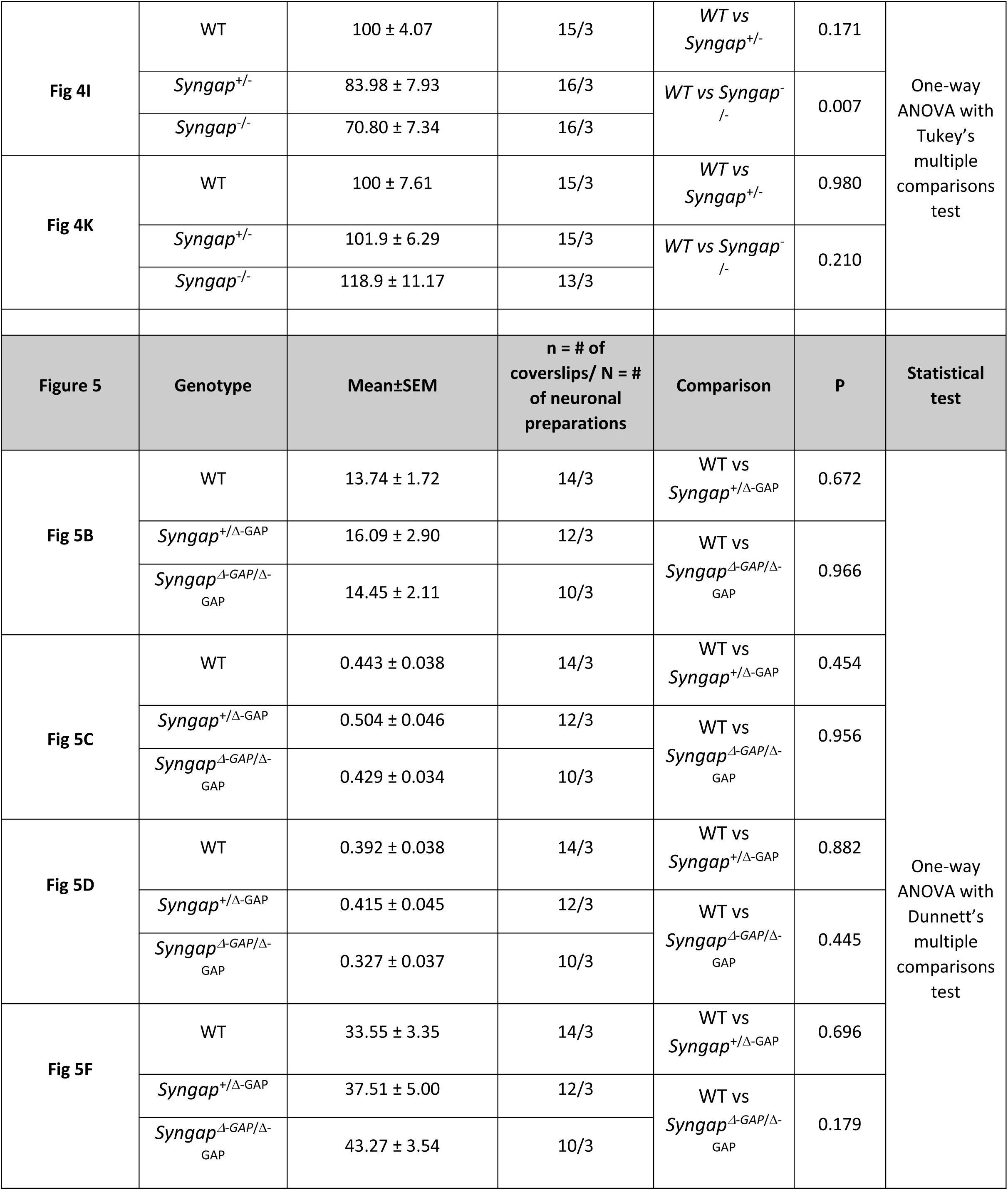
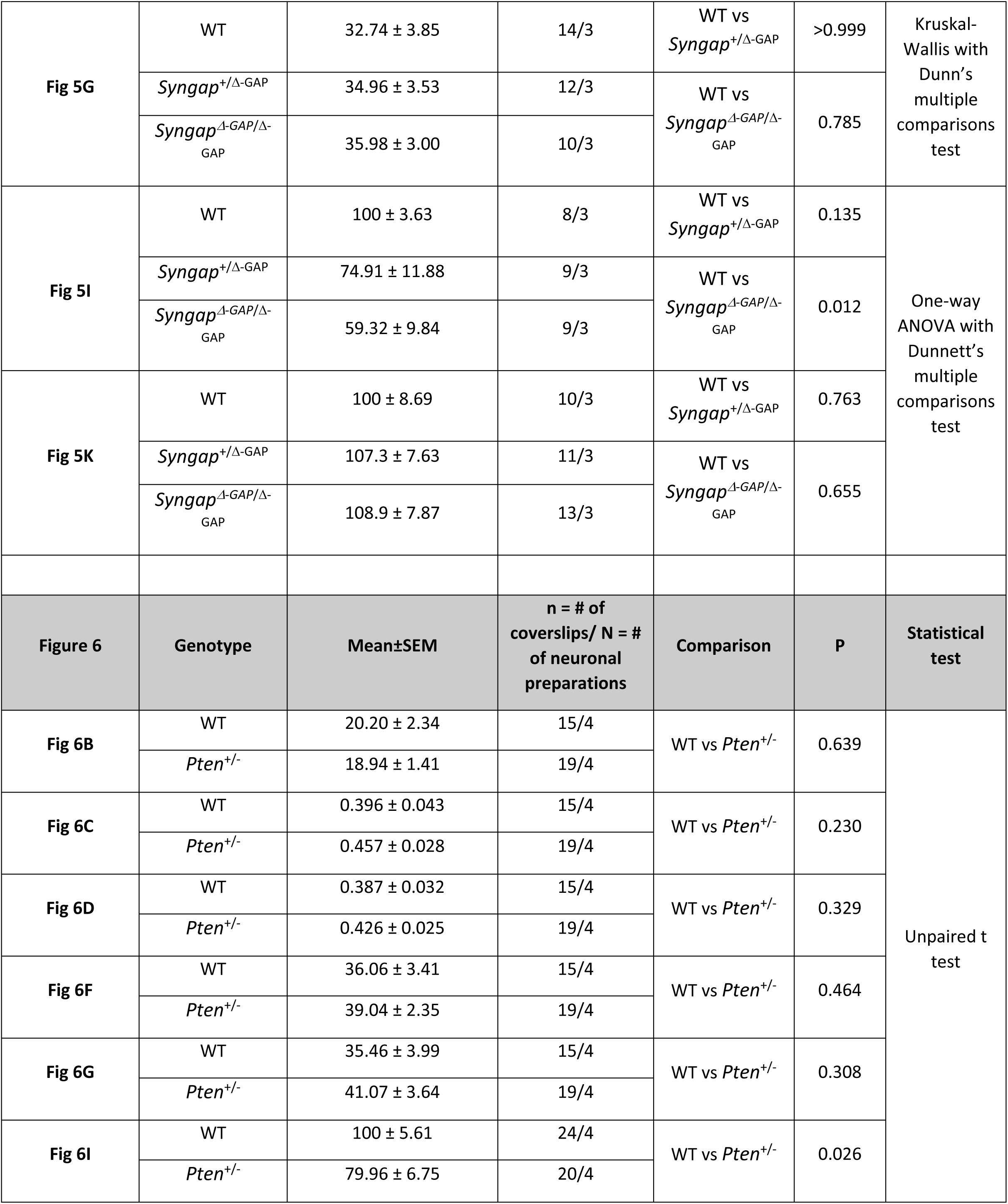
Collated table of statistical analysis.

## Acknowledgements

This work was supported by grants from the Simons Foundation (529508), Epilepsy Research UK (P2003) and the RS McDonald Fund.

